# Sleeping While Awake: The Intrusion of Neural Activity Associated with Sleep Onset in the Awake Human Brain

**DOI:** 10.1101/2020.06.04.133603

**Authors:** Stephanie Hawes, Carrie R. H. Innes, Nicholas Parsons, Sean P.A. Drummond, Karen Caeyensberghs, Richard D. Jones, Govinda R. Poudel

**Affiliations:** Mary Mackillop Institute for Health Research, Faculty of Health Sciences, Australian Catholic University, Melbourne, Australia; New Zealand Brain Research Institute, Christchurch, New Zealand; Department of Medicine, University of Otago, Christchurch, New Zealand; Turner Institute for Brain and Mental Health, Monash University, Melbourne, Australia; Cognitive Neuroscience Unit, School of Psychology, Deakins University, Melbourne, Australia; Department of Electrical and Computer Engineering, University of Canterbury, Christchurch, New Zealand; School of Psychology, Speech and Hearing, University of Canterbury, Christchurch, New Zealand

**Keywords:** Sleep Deprivation, Sleepiness, Drowsiness, Local sleep, Microsleeps

## Abstract

Sleep can intrude into the awake human brain when sleep deprived or fatigued, even while performing cognitive tasks. However, how the brain activity associated with sleep onset can co-exist with the activity associated with cognition in the awake humans remains unexplored. Here, we used simultaneous fMRI and EEG to generate fMRI activity maps associated with EEG theta (4-7 Hz) activity associated with sleep onset. We implemented a method to track these fMRI activity maps in individuals performing a cognitive task after well-rested and sleep-deprived nights. We found frequent intrusions of the fMRI maps associated with sleep-onset in the task-related fMRI data. These sleep events elicited a pattern of transient fMRI activity, which was spatially distinct from the task-related activity in the frontal and parietal areas of the brain. They were concomitant with reduced arousal as indicated by decreased pupil size and increased response time. Graph theoretical modelling showed that the activity associated with sleep onset emerges from the basal forebrain and spreads anterior-posteriorly via the brain’s structural connectome. We replicated the key findings in an independent dataset, which suggests that the approach can be reliably used in understanding the neuro-behavioural consequences of sleep and circadian disturbances in humans.

---

Brief sleep onsets can intrude into wakefulness when homeostatic sleep drive is elevated due to sleep loss, fatigue, or extended monitoring tasks. These transient sleep intrusions can be local and restricted to the momentary silencing of a few neurons (Krueger JM and G Tononi 2011; Vyazovskiy VV et al. 2011; Nir Y et al. 2017; Quercia A et al. 2018) or on a global scale; characterised by the slowing of neural activity in widespread cortical and sub-cortical regions (Boyle LN et al. 2008; Ong JL et al. 2015; Jonmohamadi Y et al. 2016; Toppi J et al. 2016; Wang C et al. 2016; Poudel GR et al. 2018). While some extreme sleep episodes are associated with behavioural signs of falling asleep such as attentional lapses (Drummond SP et al. 2005; Chee MWL et al. 2008) and slow closing of the eyelids (Poudel GR et al. 2014; Poudel GR *et al.* 2018), sleep onset can also occur without overt behavioural signs. Sleep onsets are particularly frequent in individuals who are exposed to sleep and circadian disturbances, and can acutely reduce cognitive function (Nir Y *et al.* 2017). Hence, monitoring sleep onsets while awake has major implications for understanding human behaviour in shiftwork, safety-critical operations including motor vehicle accidents, as well as circadian and sleep disorders.

Electroencephalography (EEG), pupil sizes, and response behaviour can be used to monitor overt signs of reduced arousal in humans. Any brief (3–15 s) intrusions of theta waves (4–7 Hz theta) that replace higher-frequency alpha waves (>8 Hz) on EEG recordings are considered to be microsleeps (Boyle LN *et al.* 2008). In drowsy individuals, response lapses are associated with these microsleeps (Poudel GR *et al.* 2014; Poudel GR *et al.* 2018; DiFrancesco MW et al. 2019). Eye monitoring has also been used to identify episodes of slow eye-lid closures lasting several seconds, which indicate transition to sleep whilst awake (Ong JL *et al.* 2015). However, these behavioural and neural indicators of sleep intrusions do not completely overlap (Brown RE et al. 2012; Tagliazucchi E and H Laufs 2014; Chang C et al. 2016). Not all lapses or slow-eye-closures correspond to intrusions of theta-activity on EEG, and individuals may still be responsive and report being awake even during the intrusions of early EEG-defined non rapid eye movement (NREM) sleep (Ogilvie et al. 2001). As such, arousal is an endogenous mental state with high interindividual variability in its behavioural/physiological expression, which can limit in the use of current techniques in reliably tracking sleep intrusions whilst awake.

Recent neuroimaging findings suggest that stages of arousal can manifest as dynamic changes in neural activity and connectivity as measured with blood-oxygen-level-dependent (BOLD) fMRI signal (Olbrich S et al. 2009; Tagliazucchi E and H Laufs 2014; Chang C *et al.* 2016; McAvoy MP et al. 2019; Teng J et al. 2019). Striking patterns of BOLD signal co-activation/deactivation have been associated with spontaneous slow-eye-closures or behavioural ‘microsleeps’, which can frequently occur when drowsy (Ong JL *et al.* 2015; Jonmohamadi Y *et al.* 2016; Toppi J *et al.* 2016; Wang C *et al.* 2016; Poudel GR *et al.* 2018). The fMRI amplitude fluctuations in specific brain networks can therefore be used to track moment-to-moment variations of arousal states in mammals (Chang C *et al.* 2016). Such large-scale co-activation patterns can also intrude into resting-sate BOLD fMRI data, reflecting momentary reductions in arousal during the awake resting state (Liu X et al. 2018). Notably, electrophysiological studies indicate that such large scale cortical activity during sleep is not a single synchronous event but a ‘travelling wave’, originating in specific brain regions and propagating across the cortex (Massimini M et al. 2004; Nir Y et al. 2011). For example, slow waves during NREM sleep originate in anterior cortical regions, spreading in an anteroposterior direction, covering large areas of the cortex (Massimini M *et al.* 2004). Whether the BOLD fMRI activity associated with sleep onset can be monitored in the awake brain and modelled as travelling activity, propagating via the brain’s structural connectome, has not yet been explored.

Here, using data from two studies, we developed a novel fMRI-based framework to infer sleep onsets in awake humans performing a cognitive task. First, we used simultaneous fMRI and EEG to identify typical fMRI activity patterns associated with transitions from wakefulness to early sleep. We then implemented a novel spatial regression technique to infer the intrusions of these sleep co-activation maps in awake but sleep-deprived humans performing a cognitive task. We then tested whether these sleep onsets could be inferred from fMRI alone, and if they are associated with pupillometric measures of reduced arousal. Lastly, we implemented a graph theoretical computational model to predict whether BOLD activity during inferred sleep onsets can be modelled as travelling activity, spreading via the brain’s structural connectome. The key findings were also validated using an independent fMRI dataset.

## Materials and methods

### Study 1: Simultaneous fMRI and EEG study of sleep onsets

#### Participants and protocol

Twelve healthy participants (6 female; Age: 19–27; right handed) were recruited for a 40-min daytime nap session inside a 3T MRI scanner (Siemens Skyra). Of these, ten completed the full simultaneous fMRI and EEG session. To facilitate stage 1 sleep within the MRI environment, the experiment was performed following a substantial lunch. The post-lunch circadian dip is the period of lowered arousal that occurs between 13:00 and 16:00 due to a small reduction in core body temperature, which promotes a tendency to sleep (Javierre C et al. 1996). The experiment lasted up to 2 hours, including preparation outside the MRI scanner.

#### EEG and MRI data collection

fMRI data was acquired using an echo planar imaging (EPI) acquisition that covered most of the brain (excluding cerebellum) (slice thickness = 3.3 × 3.3 × 3.3mm, TR=2.5s, TE=40ms, FA=90°, Total Scan Time = 40 mins). T_1_-weighted anatomical images (TE = 2.07ms; TR = 2.3s; field of view: 256 × 256 mm; slice thickness: 1mm) were also collected. Simultaneous EEG data was acquired using a 64-channel MR compatible EEG system (BrainProducts, Germany) as per best practice published elsewhere (Mullinger KJ et al. 2013). The EEG data were acquired at 5 kHz (Brain Vision Recorder, Brain Products, Germany). Hardware filters of 0.016–250Hz were used during the data recording (*SI Methods*).

#### EEG data analysis

We defined sleep onsets as increased power in low frequency theta activity (4–7 Hz, i.e., similar to NREM sleep stage 1) on EEG recordings. To this end, average power spectral density (PSD) in the theta (4–7 Hz) and alpha (8–13 Hz) bands were estimated for 9 occipito-parietal EEG electrodes (O1, O2, OZ, P1, P2, PZ, PO3, PO4, POZ), at each MRI repetition time (TR) (Fig. 2A). We selected these electrodes as theta waves are highest in fronto-parietal electrodes and alpha waves are highest in the occipito-parietal electrodes, during drowsiness (Lal SK and A Craig 2002). The PSD distribution at these electrodes was within the theta-alpha band (4–12 Hz) (Fig. 2B-D). EEG data were processed to remove MRI (gradient artefact) and ballistocardiogram (BCG) artefacts (SI Methods). The denoised EEG data were analyzed using a time-frequency analysis implemented in Matlab software. EEG channel data was analyzed using a moving window of 2.5s, yielding a spectrogram for each electrode via Welch’s periodogram method.

#### fMRI data analysis

fMRI data were preprocessed using FSL (FMRIB’s Software Library, www.fmrib.ox.ac.uk/fsl), Advanced Normalisation Tools (ANTs) (http://stnava.github.io/ANTs/), and custom Linux Shell and Matlab scripts (Matlab 7.6.0, R2018a, Mathworks, MA, USA). Data preprocessing steps (SI Methods) included (1) motion correction, (2) slice-time correction, (3) spatial smoothing (6-mm Gaussian kernel), and (4) high-pass filtering with a cut-off of 256s. The fMRI data was normalized to the 2 × 2 × 2mm^3^ Montreal Neurological Institute (MNI) template using linear and non-linear registration available in ANTs (SI Methods).

To identify the fMRI activity associated with EEG alpha and theta activity, the EEG alpha and theta timeseries were used as regressors in a general linear model analysis of the fMRI data (SI Methods). For each individual subject, EEG alpha, EEG theta, large motion outliers, and 6 motion parameter regressors were used in a linear regression model, which was fitted to the data in a voxel-wise manner. The regression model was estimated at the first level for each participant, generating parameter estimates maps of activity for each subject. A group-level one-sample *t*-test was performed using non-parametric statistics (FSL randomize) with 5000 permutations. The main-effects of alpha and theta on fMRI activity was considered significant at *p*<0.05 (family-wise-error corrected using cluster thresholding at z>2.3).

### Study 2: fMRI activity during logical decision making while well-rested and partially sleep deprived

#### Participants and protocol

Twenty healthy right-handed adults (10 females) aged between 20 and 37 years (M=24.9, SD=4.2) participated in the study (SI Methods). All participants had no history of psychiatric, neurological, or sleep-related disorders. Participants took part in two sessions of experiments (repeated measure design; well-rested and sleep-deprived) which occurred one week apart. These sessions were counterbalanced across participants (10 participants were well-rested in the first session and 10 were sleep-deprived in the first session). All participants were directed to maintain normal sleep behaviours during the week prior to the experimental sessions, except for the night immediately preceding their sleep-deprived testing session. In this case, time-in bed was restricted to 4 hours (3:00am–7:00am). Ethics approval for the study was obtained from the New Zealand Upper South Regional Ethics Committee.

#### MRI Data Collection

Participants in this study were imaged using a Signa HD × 3.0 T MRI Scanner (GE Medical Systems) with an 8-channel head coil. High-resolution anatomical whole-brain images were obtained using T1-weighted anatomical scans (TR = 6.5ms; TE = 2.8ms; inversion time: 400ms; field of view: 250 × 250mm; matrix: 256 × 256; slice thickness: 1mm). Functional images were obtained using EPI (TR = 2.5 s; number of repetitions = 293 TE = 35ms; field of view: 220 × 220mm; number of slices: 37; slice thickness: 4.5 mm; matrix: 64 × 64).

#### Game of set task

The participants performed a two-choice logical decision-making task, based on a ‘game-of-set’, inside the MRI scanner. In this task, the participants were presented with a set of cards on a screen and a 2-choice decision to make. The participants had to select whether the set cards were part of a set (‘yes’) or not a set (‘no’). The game-of-set rule is such that if the two cards were the same for any feature (colour, number, symbol, or shading) and one was not, then the three cards were not a set, otherwise they were a set. Participants were trained on the rules of game of set by providing detailed instruction (SI Methods). The participants had to press left (‘yes’) or right (‘no’) button using an MRI compatible button box (Fig. S5). Experimental stimuli were presented on a screen for 5.0s, followed by a fixation cross for 2.5–10 s (Mean = 4.0, jittered). Each participant completed three runs of 6-min duration. Each run consisted of 40 experimental stimuli and 40 fixation crosses (total trials =120 in 3 runs). Performance was measured through response time. The participants were provided training on the task prior to the start of the scanning session.

#### Pupillometric data collection

Eye-video data were captured whilst participants completed the task. Right-eye movement was recorded on a Visible Eye™ system (Avotec Inc., Stuart, FL) mounted on the head-coil of the MRI scanner. Custom-built video recording software and a video-capture card was used to record eye-videos onto a computer at 25 frames/s (350 × 280 px). The eye-video data was processed using the starburst algorithm in Matlab software to measure the participant’s pupil size from the dark-pupil infrared illuminated eye videos (Dongheng Li et al. 2005). Relative change in pupil size is a good indicator of changes in arousal (Chang C *et al.* 2016; Liu X *et al.* 2018); hence we estimated the time-course of pupil size at the onset of inferred sleep events by calculating baseline corrected time-locked pupil size.

#### Preprocessing of fMRI data from Study 2

The fMRI data from study was preprocessed and normalized as per the details described for Study 1.

#### A method for detecting sleep onsets in the awake human brain

The fMRI activity pattern associated with EEG theta (Fig. 2E, Fig. S1) is archetypical of reduced in arousal associated with early sleep-like behaviour in both humans and macaques (Chang C *et al.* 2016; Liu X *et al.* 2018; Stevner ABA et al. 2019). Hence, we postulated that by monitoring intrusions of the fMRI activity map associated with EEG-theta activity, we may be able to infer sleep onsets in the fMRI data from the study 2 without the need for simultaneous EEG recordings. Thus, we implemented a method to track EEG-theta fMRI activity patterns in the fMRI dataset from the Study 2 (Overview provided in Fig. 1). First, we mapped average theta-related fMRI activity (Fig. 2, Fig. S1) in 82 brain regions by using the cortical and sub-cortical parcellations from the Desikan-Killiany atlas, resulting in a theta-activity vector (82 × 1). A rolling window regression model was then employed to identify any transient fMRI activity in pre-processed fMRI signal (motion corrected, normalised, task and noise related activity regressed out). We modelled transient fMRI activity as a typical haemodynamic response function with span of 32.5s (13 TR). At each time-point, this model was fit to the data using a general linear model. By using a ‘rolling regression’, we identified parameter estimates of the fit between the transient fMRI model and the denoised fMRI data in each of the 82 regions, resulting in 82 × 1 vectors of parameter estimates at each time point. These parameter estimates were then correlated (Pearson’s correlation) with the 82 × 1 vector of fMRI activity associated with EEG-theta activity, which provided as estimation of how well the transient activity at each time-point represents sleep-like activity. Any time-points where the correlation was significantly high (*p*<0.05, corrected) and positive (i.e., *r* > 0) were considered to be due to sleep onsets (inferred sleep onsets).

**Fig 1.**
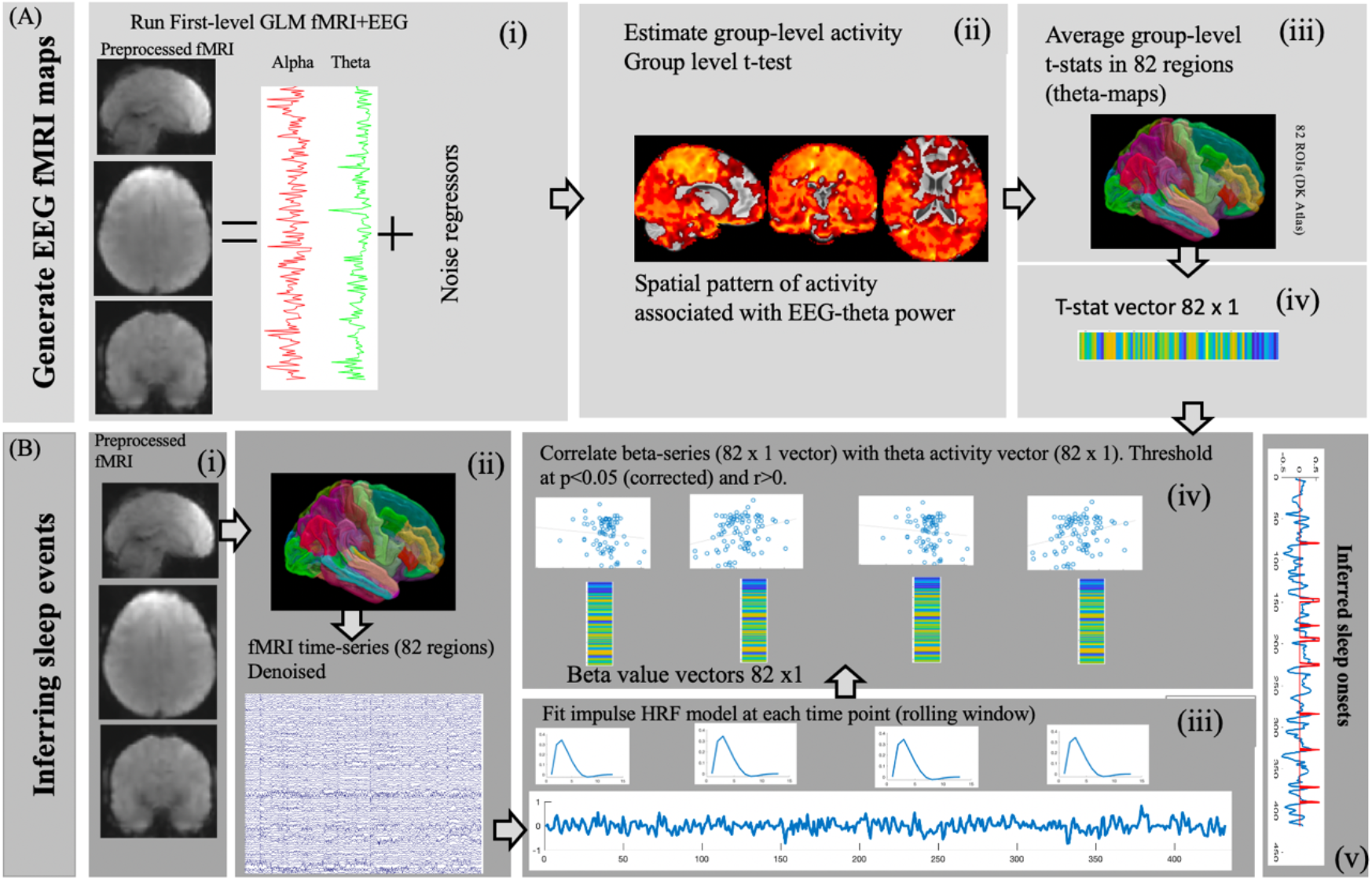
Graphical overview of the method implemented for detecting sleep onsets from fMRI data. (A i-ii) Simultaneous fMRI and EEG data were analyzed using a GLM to generate a spatial template of fMRI activity associated with power spectral density in EEG-theta. (A iii) Average BOLD fMRI activity (t-statistics) associated with EEG theta was estimated for 82 regions as per the Desikan-Killiany atlas. (A iv) This process generated a vector of 82 ×1 in size corresponding to the EEG-theta related BOLD signal in 82 regions. (B i) The preprocessed and normalized fMRI data from Study 2 sleep was processed to remove noise and task-related activity. (B ii) BOLD time-courses from 82 regions (as per DK atlas) were extracted. (B iii) For each time-course, a rolling-window general linear model was run with a typical impulse haemodynamic response function as a predictor, which is run for each TR. (B iv) The beta-values of the fit between impulse response and fMRI signal for each region generates 82 × 1 vector at each time point, which is correlated against the EEG-theta vector (from A iv). (B v). Any time-points where the correlation was significant (*p*<0.05, corrected) and positive (i.e., *r* > 0) were considered to be attributed to sleep onsets.

**Fig. 2:**
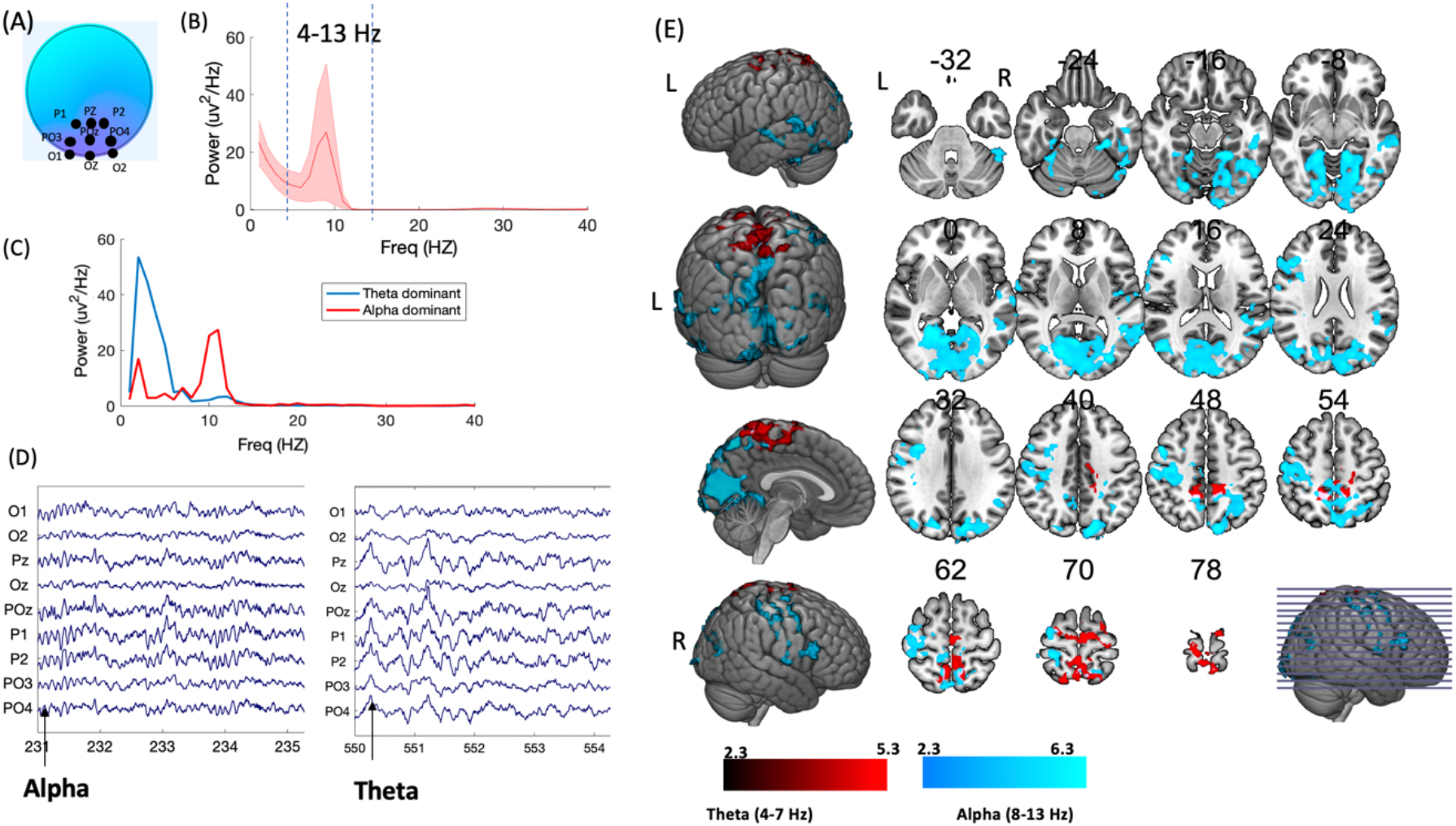
Simultaneous fMRI and EEG derived spatial pattern associated with early (stage 1) sleep. (A) EEG data from electrodes located at the occipito-parietal areas were used to identify fluctuations in alpha (8–13 Hz) and theta (4–7 Hz) power spectral density. (B) A plot showing that the average power spectral density at these electrodes was within the theta-alpha band (4–13 Hz). Shaded areas denote standard error of mean (SEM). (C-D) Examples of intrusions of alpha and theta activity in resting-EEG. (E) fMRI activity pattern associated with theta and alpha power in EEG. A positive correlation between EEG theta and fMRI signal is shown in red. A Negative association between fMRI signal and EEG alpha waves is shown in blue. Correlations were significant at the cluster-corrected level *p*<0.05 (cluster forming threshold of z=2.3). The fMRI activity are overlaid on a MNI brain with neurological orientation.

#### Voxel-wise analysis of fMRI activity associated with task and inferred sleep events

To tease apart the spatial distribution of brain activity associated with task performance and sleep onsets, we further analysed the fMRI data from study 2 using the multi-level voxel-wise modelling of fMRI data in FSL. At the subject level, the fMRI data from each participant were analyzed using a first-level general linear model, which included predictors for (1) task-related regressors (120 trials across 3 runs) modelled as epochs of duration modulated by response time, (2) sleep onsets modelled as impulse activity convolved with a double-gamma haemodynamic response function, (3) response errors modelling the time points when subjects failed to respond during the task, (4) six motion parameters, and (5) large motion outliers (from *fsl_motion_outliers* using the *dvars* option). For each subject, the main effects of task and inferred sleep events were estimated using first-level contrasts representing average activity. For second-level group analysis, a non-parametric approach was used to estimate group-level significance of the first-level parameter estimates. A group-level t-test was performed using non-parametric statistics (FSL Randomize) and 5000 permutations. The main-effect of tasks were considered significant at *p*<0.05 (voxel-level family-wise-error corrected). A paired-test model was then used to estimate difference in task-related and sleep onset related fMRI activity between rested and sleep-deprived sessions. The difference was considered to be significant at *p*<0.05 (voxel-wise FWE corrected). The main-effect of sleep onset was considered significant at *p*<0.05 (cluster corrected, z-threshold >4).

#### The impact of sleep onsets on task-related activity and connectivity

To investigate the potential impact of sleep onsets on task-related functional activity and connectivity, we analysed task-related activity and connectivity in six key bilateral brain regions associated with the task (using main-effect of task during rested session only). These regions were chosen based on their involvement in decision-making (Poudel GR et al. 2017). These included the bilateral prefrontal, anterior cingulate, inferior parietal, putamen, thalamus, and insula. Average task-related activity in each individual was extracted using spherical masks (10 mm radius) centered on the local-maxima of activity within each region. The average task-related activity was then correlated (Pearson’s r) with the total number of inferred sleep events in both rested and sleep-deprived sessions. Beta-series correlation was used to estimate task-specific functional connectivity between the six regions (Rissman J et al. 2004). This method implements separate predictors to model task-related activity using general linear model (GLM). The resulting parameter estimates (beta values) for each brain regions were correlated to derive a task-related functional connectivity matrix. The association between functional connectivity and sleep intrusions were then estimated by using Pearson’s correlation and considered significant at *p*<0.05 (FWE corrected).

#### Computational modelling of the spread of sleep-related neural activity

We modelled the spread of sleep-related activity in the brain as a passive, diffusive flow of activity via brain’s structural connectome (Poudel GR et al. 2019; Poudel GR et al. 2020). We used the structural connectivity of a normative brain that has been made publicly available (IIT Human Brain Atlas V5.0 https://www.nitrc.org/projects/iit). The structural connectome was defined for the 82 cortical- and sub-cortical regions as per the Desikan-Killiany atlas. This undirected brain network graph can be represented as **G** = (**ϖ, ε**), where **v** is the set of brain parcels (nodes) given by **v**=(v_*1*_, v_*2*_, … , v_*n*_) and ε is the set of connections between **v**_*i*_ and **v**_*j*_ (edge) given by **ε** = (v_*i*_, v_*j*_). The spread model treats the edge (v_*i*_, v_*j*_) as a conduit that connects nodes v_*i*_ and v_*j*_, and the spread of activity at time *t* can be modelled as:

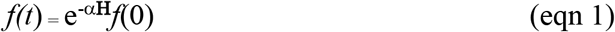

where *f(t*) denotes the vector consisting of the amount of diffusion of pathology at node v_*i*_ at time *t*, beginning from an initial distribution of pathology given by *f(0*) at time zero. **H** is the graph Laplacian defined as the difference between degree matrix and adjacency matrix. Alpha (α) is the spread constant (assumed to be 1 in our analyses).

The activity at a node at time t *f(t*) can be estimated as:

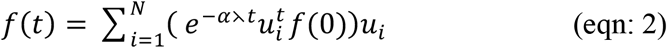

where *U*=[u_1_, u_2_, u_3_, …u_n_] represent eigenvectors of the Laplacian matrix.

To assess whether the initiating spread of brain activity in any specific region of the brain is most predictive of patterns of sleep-related activity, we simulated the process of spread on a normative connectome by repeatedly initiating the spread from all brain regions within the Desikan-Killiany atlas. For each region *i*, the network model is used to estimate the spread at all other regions at time *t* = 1−20, with initial condition *f(*0) set as a unit vector with 1 at the *i*^th^ location and 0 elsewhere. This process generated vectors with 82 elements at each time point. The predicted activity vector at all time points, *f(*t), were correlated against measured group-level activity (*t*-value) using Pearson’s correlation coefficient. For each seed region *i*, we identified the maximum correlation value, which was used as a measure of the likelihood of the region being the putative seed of the spread (Fig. S4).

To test for the specificity of our findings, we evaluated the model against two null models. First, we randomly scrambled the group-level activity (*t*-stats) of the neural activity vector and simulated the spread model on a true connectome. Second, we ran the spread model on 1000 random connectome with preserved degree distribution of the original connectome. These two analyses allowed us to infer whether the true prediction was significantly greater than a prediction using random network.

#### Validation of the key finding using an independent dataset

To test the robustness of our findings and validate our approach for detecting transient neural activity associated with sleep onsets in fMRI data, we also analyzed another independent dataset (N=56) of partially sleep-deprived participants from the Open neuro database (ds000201). The data were processed using identical steps outlined in previous steps. Further details on the processing of this data is provided in the Supplementary Information.

## Results

### Spatial pattern of fMRI activity associated with sleep onset

A positive association (p<0.05, cluster corrected at z>2.3) between EEG theta and fMRI signal was observed in the bilateral precuneus, superior parietal lobule, precentral gyrus, postcentral gyrus, and superior frontal gyrus (Fig. 2E, Table S1, Fig. S1). A negative association (p<0.05, cluster corrected at z>2.3) was observed between EEG alpha and fMRI activity in wide-spread cortical regions (Fig. 2E, Table S1, Fig. S1) including the bilateral precuneus, superior parietal, precentral, postcentral, lateral occipital, middle frontal, and middle temporal cortices. A negative association was also observed in deeper cortical regions including the parahippocampus and insula.

### Frequent intrusions of sleep during a cognitive task performed while rested and sleep-deprived

In the study 2, the total number of the sleep onsets, detected using novel framework, ranged from 17 to 63 (Mean: 35, SD: 14) in rested sessions and 20 to 74 (Mean: 42, SD: 15) in sleep-deprived sessions (paired t-test, *t*(18) = 1.6, *p* = 0.1) (Fig. 3A, Fig. S2). There was a significant difference in average pupil size during the sleep onsets detected at rested and sleep-deprived sessions (*t*(18) = 3.44, *p* = 0.002), with on-average a 20% reduction in pupil size after sleep-deprivation (Fig. 3B). The decision trials which coincided with the sleep onsets had longer average response time (RT) compared to the other trials (*t*(18) = 2.21, *p* = 0.04) when in the sleep deprived session. There was a positive correlation between total number of sleep events and mean RT in the sleep-deprived session (*r*=0.53, *p*=0.02) but not in the rested session (Fig. 3C).

**Fig. 3:**
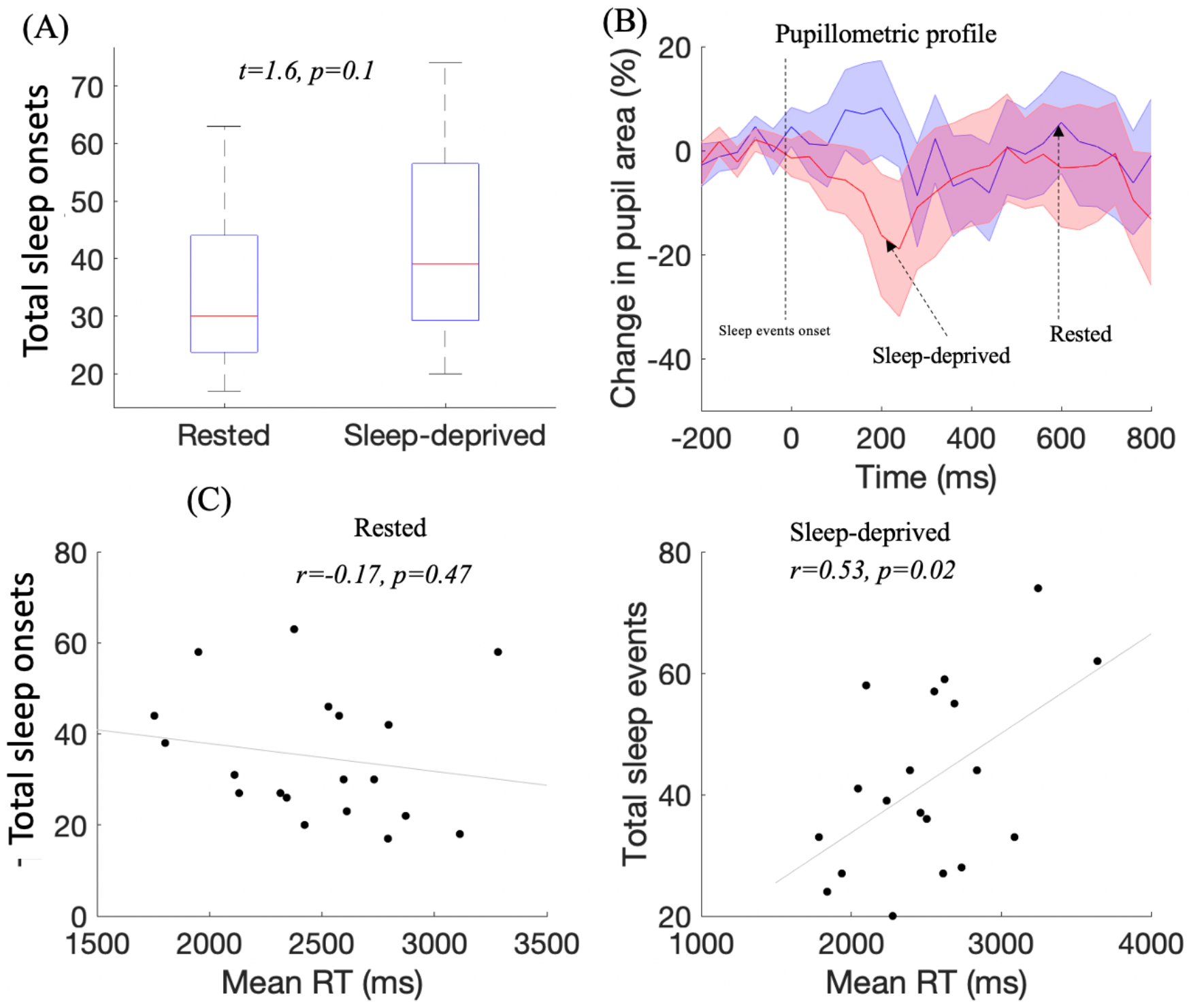
The pupillometric and behavioural characteristics associated with the sleep onsets detected from fMRI data. (A) Boxplot of mean and SD of the total number of sleep onsets in rested and sleep-deprived conditions. There was no significant difference in total number of sleep onsets. (B) The plots of changes in pupil area from baseline during sleep onsets. Shaded error bars represent SEM. There was on average a 20% reduction in pupil size during inferred sleep events after sleep-deprivation. (C) Pearson’s correlation between total number of sleep onsets and average reaction time (mean RT) during the task. There was a significant correlation during sleep-deprived session but not while well-rested.

### Spatially distinct fMRI activity during the task and sleep onsets

The spatial patterns of fMRI activity associated with task and sleep onsets in rested and sleep-deprived sessions are provided in Fig. 4 A, B (Table S2). Significant task-fMRI activity (*p*<0.05, FWE-corrected) was observed in the bilateral prefrontal (inferior/middle frontal), motor (precentral/postcentral), parietal (superior parietal), anterior and posterior cingulate, and bilateral insula cortices. Increased sub cortical activity was also observed in the bilateral thalamus and striatum. There was no significant voxel-wise difference in task-related brain activity between rested and sleep-deprived sessions.

**Fig. 4:**
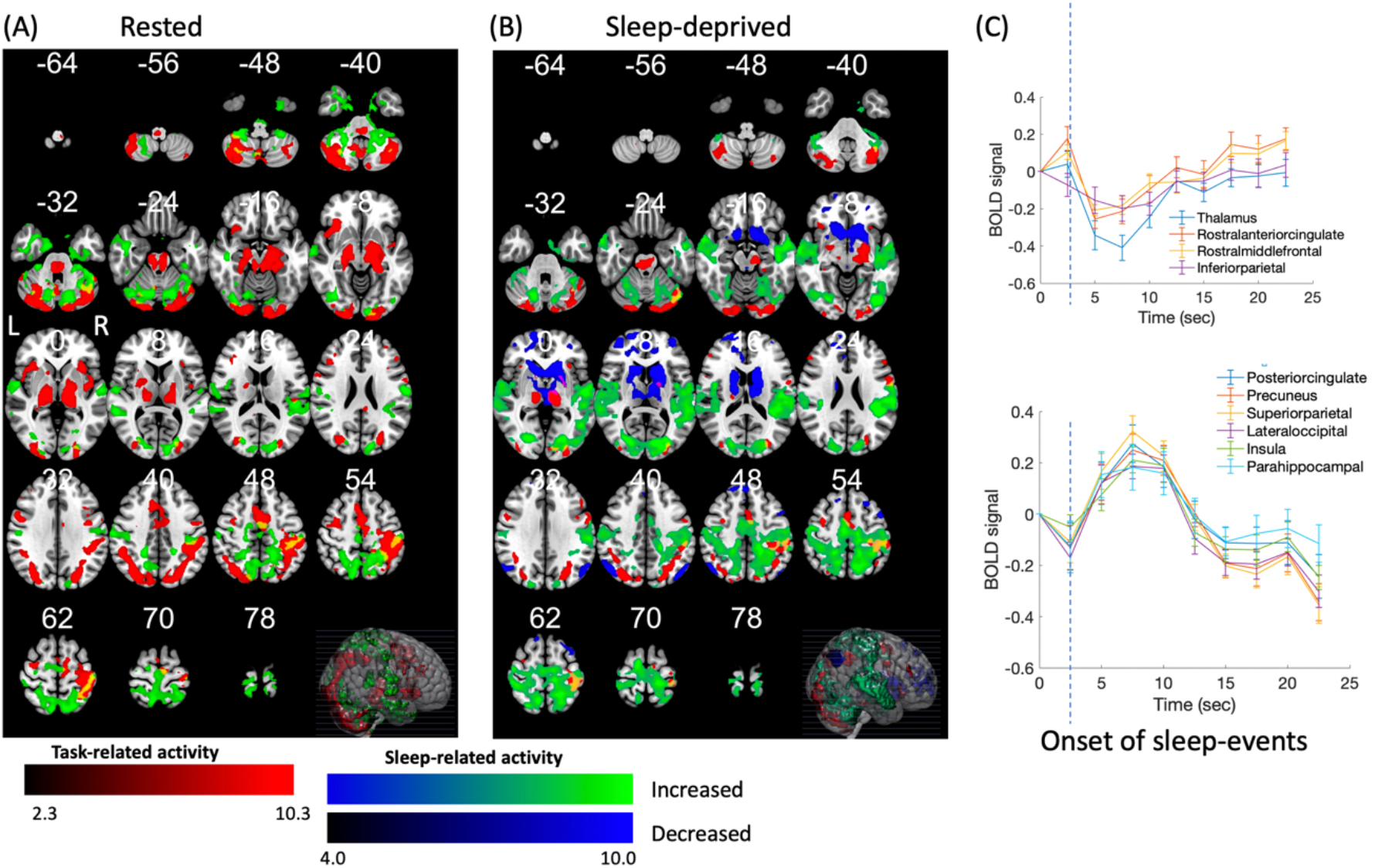
Whole-brain fMRI activity patterns associated with the decision-making task and inferred sleep events. (A) The pattern of task-related activation (red, *p*<0.05 FWE-corrected) overlaid with sleep-onset related activation (green, *p*<0.05, cluster-corrected) observed during the rested session. (B) The pattern of task-related activation (red, *p*<0.05 FWE-corrected), sleep-onset related activation (green, *p*<0.05, cluster-corrected), and sleep-onset related deactivation (blue, *p*<0.05, cluster-corrected) observed during the sleep-deprived session. Any overlap in task-related and sleep-related activation is rendered in yellow. (C) Baseline corrected time-courses (and SEM) of fMRI signal time-locked to the sleep events (marked by dotted line) in several brain regions, showing synchronous activation/deactivation patterns in the wide-spread cortical and sub-cortical regions

Whereas, fMRI activity during inferred sleep onsets (*p*<0.05, cluster-corrected, Z≥4.0) was observed in the bilateral visual (occipital pole, cuneus, lingual gyri), auditory (superior temporal gyri, Heschl’s gyri), primary and secondary somatosensory (postcentral gyri, parietal operculum, superior parietal lobule, and insular cortex) areas, the primary motor (precentral gyri) and supplementary motor areas, default-mode areas including in the bilateral precuneus and angular gyri and limbic areas encompassing the bilateral parahippocampal regions (Fig 4 A,B). Decreased activity was observed in sub cortical areas including the bilateral thalamus, caudate, and putamen, and cortical areas such as the rostral anterior cingulate gyrus (basal forebrain), rostral middle-frontal gyrus, and lateral inferior parietal areas. These deactivations were only significant in the sleep-deprived session. The time-courses of fMRI activations and deactivations at the onset of sleep-events in a number of cortical and sub-cortical regions are shown on Fig. 4C. Time-courses demonstrated a clear pattern of synchronous activations in cortical regions. Whereas, deactivation appeared to be strongest in the bilateral thalamus. There was also some overlap between the task-related and sleep-related networks, particularly in the motor/association cortex. This overlap was observed in the left precentral gyrus, supplementary motor area, left superior parietal cortex, and bilateral lateral occipital cortices (Fig S3).

### The impact of sleep onsets on task-related activity and connectivity

There was a significant negative correlation between total sleep events and level of fMRI activity in the left thalamus (*r*=−.53, *p*=0.02) and bilateral putamen (*r*=−.58, *p*=0.009) during the sleep-deprived condition (Fig. 5B). Other regions of interest did not show any association either in the well-rested or sleep-deprived conditions. Task-related functional connectivity between the left ACC and right putamen, the left ACC and right thalamus, the left ACC and right dorsolateral prefrontal cortex (DLPFC), and the right DLPFC and right putamen were significantly positively correlated (p<0.05) with total sleep events in the sleep-deprived session (Fig 5A, C). There was no significant correlation however between functional connectivity and total sleep events in the rested session (Fig. S4).

**Fig. 5:**
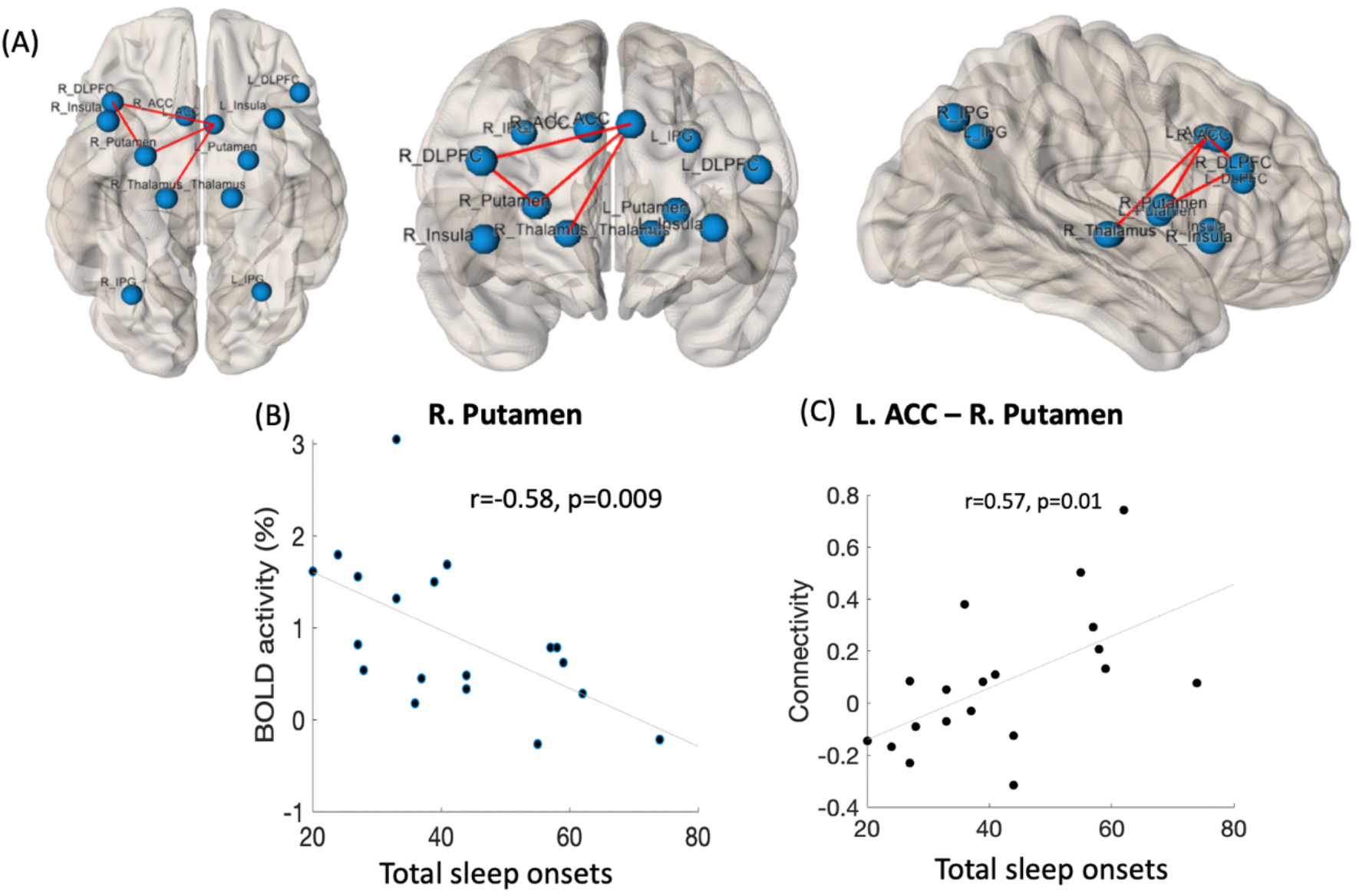
Visualization of the association between task-related activity and functional connectivity and intrusions of sleep onsets. (A) The cortical and sub-cortical regions of interest including right and left DLPFC, right and left thalamus, right and left ACC, right and left insula, right and left inferior parietal lobule are visualized on a surface brain. The functional connections showing significant association between functional connectivity and total sleep-onsets are visualized as red lines. Significant correlations were observed between total number of sleep onsets and task-related fMRI activity in the left thalamus, right putamen, and left putamen. (B) Example scatterplots showing significant association between number of sleep onsets and fMRI activity/functional connectivity in the right putamen and left ACC – right putamen functional connection. Remaining scatterplots are provided in the supplementary data (Fig S4).

### Modelling the spread of neural activity during sleep-like intrusions

The network spread generated predictions of neural activity over time, which was correlated with the actual neural activity pattern associated with inferred sleep intrusions (i.e. Fig 4). We identified that seeding the neural deactivation from the medial orbitofrontal region (basal forebrain) provided the best fit between predicted and measured activity at all time points (Fig. 6 A, B). This association was significant for the sleep-onset related activity from both rested and sleep-deprived sessions (Fig. 6 C)

**Fig. 6:**
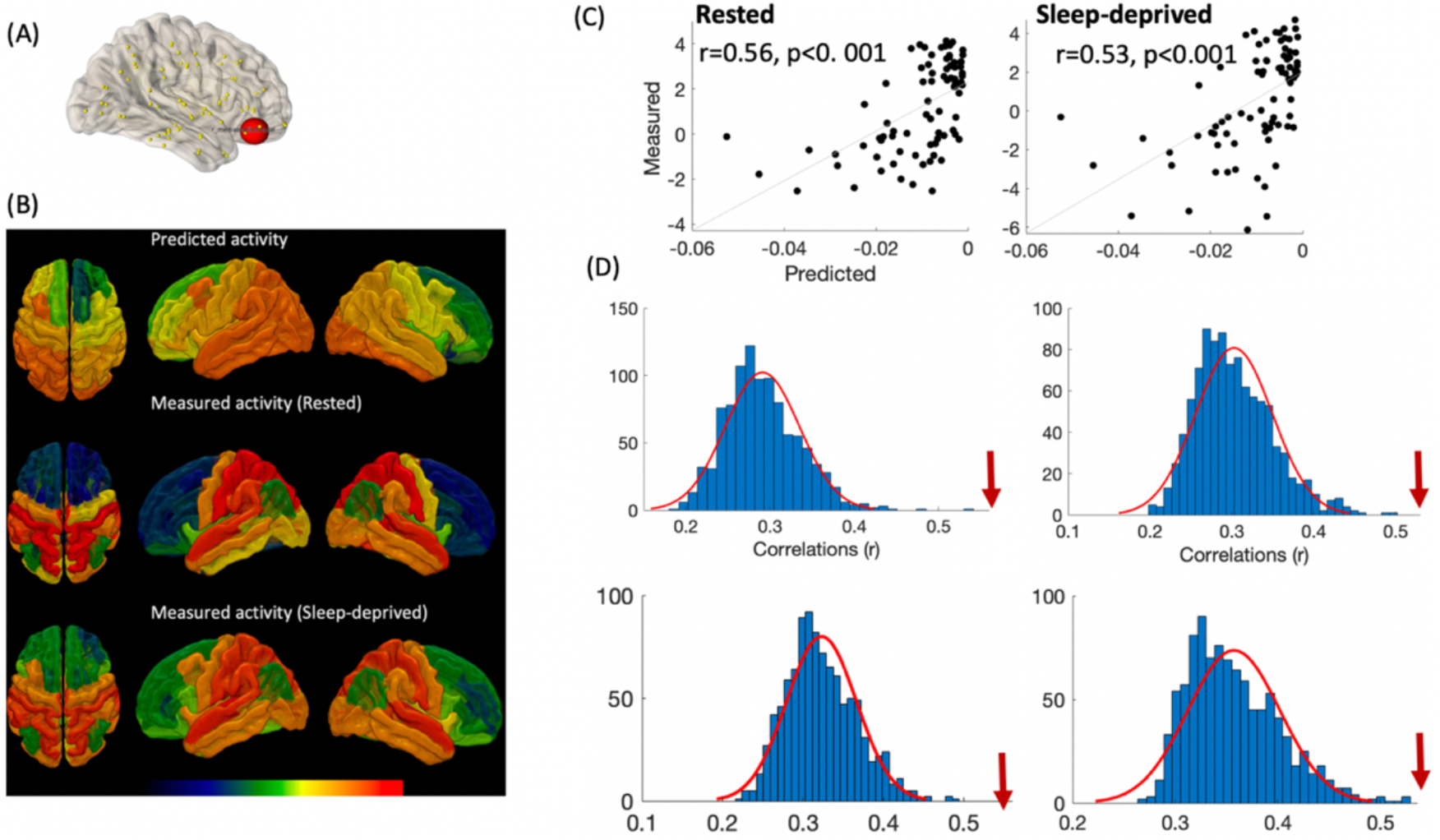
Trans-connectome spread of activity determines the pattern of sleep-related BOLD activity in the brain during intrusions of sleep in the awake brain. (A) The basal forebrain seed was the best predictor of BOLD activity during sleep-like intrusions. (B) The spatial pattern of predicted and measured activity when seeding the spread of deactivation is shown overlaid on the Desikan-Killiany atlas. (C) The correlation between predicted and measured activity was highest for the basal forebrain seed in both rested and sleep-deprived sessions. (D) Histograms of correlations when the model was evaluated against a randomly permutated activity pattern (100 times, top panel) and null networks (1000 times, buttom panel). The correlation achieved from a true model (red arrow) is well outside the 95% confidence interval of correlations achieved by random models.

The findings are specific and are significantly higher compared to achieved using random networks and random neural activity pattern. The distribution of prediction (correlation values) over 1000 scrambled matrices is shown in Fig. 5D (top panel). The random model’s correlation values are much lower than the correlation of 0.56 (0.53 in rested) achieved by the true model and are outside the 95% confidence interval, or *p* < 0.05. Hence, the reported prediction of neural activity pattern seeded from the basal forebrain may not be explained by chance.

### Replication of key findings using an independent dataset

In the independent validation dataset, the inference method identified frequent intrusions of sleep-like events (Mean: 23, SD: 8) in the partially sleep-deprived participants. The events were used in a whole-brain GLM analysis of the fMRI data, which identified co-activation and de-activation patterns similar to the ones observed during sleep-like activity in our study (see SI; Fig 7 A, C). The individuals who reported to be very sleepy during the scan (Using median split of KSS scores, median KSS>7.5) showed significantly greater activity in the visual cortex (unpaired t-test, p<0.05, cluster corrected at z>2.3, local maxima MNI: −12, −74, −6) compared to alert individuals.

## Discussion

The competitive neural systems underpinning wakefulness and sleep can be unstable in individuals who are vulnerable to sleep and circadian disturbances. This instability has been linked to brief intrusions of local and global sleep-like states during wakefulness (Krueger JM and G Tononi 2011; Vyazovskiy VV *et al.* 2011) (Olbrich S *et al.* 2009; Nir Y *et al.* 2011; Ong JL *et al.* 2015; Poudel GR *et al.* 2018). However, there are still many unanswered questions regarding how the states of sleep and wakefulness can co-exist in humans and how these states manifest in human behaviour. We have developed a method and a model to track the ebb and flow of neural activity associated with sleep onsets in awake and cognitively active humans. Our results can be summarized into three main findings; (1) The neural activity associated with sleep onsets, identified using simultaneous fMRI and EEG, can frequently intrude into awake humans. (2) These inferred sleep onsets are associated with pupillometric markers of reduced arousal and slowed response times, particularly following sleep-deprivation. (3) Sleep onsets are associated with a transient pattern of activation and de-activation in brain networks distinct from co-existing task-related brain regions. (4) Graph theoretical modelling showed that a model of spread of neural activity from the basal forebrain, via the structural connectome, can predict the measured pattern of neural activity during sleep onsets, suggesting that the basal forebrain acts an epicentre for the propagation of neural inhibition during the transition into sleep.

### EEG-theta activity at sleep onset is associated with an increase in fMRI activity

The transition from wakefulness to sleep is considered to be a state characterised by the slowing down of neural activity, the disappearance of higher-frequency waves (>8 Hz) and appearance of slower waves (1–7 Hz) on EEG recordings. However, recent neuroimaging findings suggest that the sleep onset process is dynamic and can have both transient increases and decreases in regionally-specific brain activity (Dang-Vu TT et al. 2010; Krueger JM and G Tononi 2011; Quercia A *et al.* 2018). Consistent with this view, we found that early sleep characterised by increased power in EEG theta waves is associated with increased BOLD activity in the posterior parietal areas of the brain. Whereas alert wakefulness indicated by increased power in EEG alpha waves is associated with decreased activity in the occipital, parietal, frontal, and limbic areas of the brain. When relaxed with eyes-closed, waxing and waning of alpha activity can occur due to a drift into either the drowsy or alert state (Laufs H et al. 2003). The transition towards a more alert/attentive state is associated with increased fronto-parietal activity (Laufs H *et al.* 2003). In contrast, transition to drowsiness manifests as increased occipito-parietal activity (Laufs H *et al.* 2003). By modelling EEG theta-related activity in each individual, we were able to isolate and replicate regionally specific increased activity during increased EEG theta activity; a pattern of activity typically associated with lower arousal and early sleep in humans (Laufs H *et al.* 2003; Kaufmann C et al. 2006; Laufs H et al. 2007; Brodbeck V et al. 2012; Tagliazucchi E and H Laufs 2014) and other mammals (Chang C *et al.* 2016).

### Sleep onsets can intrude in both well-rested and sleep-deprived individuals

Emerging evidence suggests that behavioural states of sleep and wakefulness are not discrete mental states but may overlap and even co-exist. A sleep-like neural activity pattern (i.e., momentary neural silencing) can intrude into an awake and active brain in response to a prolonged period of activity or increased homeostatic sleep drive (Krueger JM and G Tononi 2011; Quercia A *et al.* 2018). Intrusions of brief sleep-like episodes have been reported in individuals performing monotonous monitoring tasks for an extended time (Peiris MTR et al. 2006; Poudel GR *et al.* 2014), even while well-rested. In this study however, we detected frequent intrusions of neural activity pattern associated with early sleep, identified using an independent recording of simultaneous EEG and fMRI, in awake individuals performing a cognitive task. Notably, such intrusions were not just limited to the sleep-deprived condition but were also frequent while well-rested (Peiris MTR *et al.* 2006; Poudel GR *et al.* 2014) meaning more individuals are at risk of these events than once thought.

In the current study, we extended on the research of (Chang C *et al.* 2016), who used a similar approach for tracking the tonic level of arousal in monkeys using fMRI signal. However, our approach differs substantially by allowing for tagging each intrusion of sleep-like neural activity as an sleep-onset in both rested and sleep-deprived brain. Notably, there was no difference in the frequency of inferred sleep events between rested and sleep-deprived conditions. However, there were differences in behavioural features. The sleep events in sleep-deprived sessions were associated with reduced pupil size and an increased overall response times during the decision-making trials. These behavioural differences may be explained by differences in the magnitude of sleep deprivation, and subsequent drive to sleep. Extreme sleep intrusions, which are more frequent following sleep loss, manifest as momentary slow closures of eye-lids accompanied by responses lapses (Wang C *et al.* 2016; Poudel GR *et al.* 2018; Teng J *et al.* 2019). However, local neuronal assemblies may momentarily go to use-dependent sleep, even in rested individuals without any overt sleep pressure (Krueger JM and G Tononi 2011). This is also demonstrated in studies showing that time-on-task errors can appear during monotonous tasks, without overt behavioural signs of sleep (Thiffault P and J Bergeron 2003). Transient intrusions of behavioural lapses have been documented in previous studies of performance extended monitoring tasks, even when well-rested (Peiris MTR *et al.* 2006; Huang RS et al. 2008; Poudel GR *et al.* 2014). Sustained performance on a task is therefore sufficient to induce local silencing of neuronal activity, resulting in sleep-like behaviour in humans (Quercia A *et al.* 2018). Taken together, the findings suggest that sleep-like mental states can intrude in both rested and sleep-deprived individuals performing cognitive tasks.

### The brain activity associated with sleep onsets and cognitive functioning can co-exist in spatially distinct brain regions

The sleep-onset process is regulated by mutually inhibitory pathways originating in the brainstem and hypothalamus, which act against each other, oscillating or switching on and off. One important aspect of this sleep-switch model is that sub-cortical sleep switching may be communicated with cortical brain networks, conveying neural information to the cortex and back via the thalamus and ascending arousal system. Hence, individuals with elevated sleep-drive show both increased and decreased activity in the cortical networks depending on whether they are sleepy or alert (Poudel GR et al. 2012; Toppi J *et al.* 2016). Consistent with this network model of wake-sleep regulation, voxelwise analysis of our fMRI revealed that distinct brain networks may diverge at the edge of wakefulness and sleep. Specifically, while the somatosensory and limbic areas of the brain transiently increase in activity, the thalamus and the ventral prefrontal cortices are transiently deactivated. This transient decrease in the thalamic and rostral prefrontal cortex may reflect the withdrawal of the ascending arousal system during transition into sleep (Xu M et al. 2015). Local intra-cortical recordings suggest that the thalamus is associated with decreased activity earlier than the cortex during transition to NREM sleep (Magnin M et al. 2010). Neuroimaging studies have corroborated these findings and suggest transient decreased activity in the thalamus during microsleeps (Poudel GR *et al.* 2014) and spontaneous slowed eye-closures associated with drowsiness (Ong JL *et al.* 2015; Wang C *et al.* 2016).

Furthermore, consistent with our findings, the brain also shows increased activity in somatosensory and associative brain networks at the onset of sleep and drowsiness (Horovitz SG et al. 2008; Olbrich S *et al.* 2009; Ong JL *et al.* 2015; Wang C *et al.* 2016). Such co-activation of neural activity during a hypoactive behavioural state has been attributed to rich endogenous mental activity that can occur at the sleep onset in humans (Ong JL *et al.* 2015). However, a pattern strikingly similar to ours has also been demonstrated in macaques during wake-sleep dip transitions (Chang C *et al.* 2016), suggesting that opposing neural signaling between the thalamus and cortex may be at play during sleep onset. Such trans-network divergence in neural activity may also reflect mutually inhibitory dynamics akin to the sub-cortical switching associated with the wake-sleep transition (Saper CB et al. 2005). Notably, despite the frequent sleep onsets intrusions in the brain, we found that the task-related brain networks were robustly activated in both rested and sleep-deprived sessions. That is, the task activated brain regions expected to be involved in attention and working memory (frontal and parietal regions), motor (left primary motor), decision-making (ACC, insula), and alertness/arousal (bilateral thalamus) processes. Although there was some overlap between sleep-related activation and task-related pattern, particularly in the association cortex, the pattern of deactivation was only observed in sleep-like intrusions. Furthermore, sleep deprivation did not significantly change task-related BOLD activity. However, there was a moderate association between the number of sleep onsets and task-related BOLD activity (in bilateral putamen and thalamus) and connectivity (putamen-ACC), but only when individuals were sleep deprived. Taken together, these findings suggest that distinct sleep-like brain states can co-occur with the brain-states associated with the awake and the alert brain.

### Trans-connectome spread of the brain activity during sleep onset

Although state transitions from wakefulness to sleep appear to be spontaneous, high density EEG studies have shown that underpinning neural oscillations are travelling waves which propagate anterior-posteriorly throughout the cortex (Massimini M *et al.* 2004). In the current study, we modelled the propagation of neural activity as a linear diffusion process across the brain’s structural connectome. This model suggests that the basal forebrain regions may act as the epicentre of deactivation, as the spread from the basal forebrain best predicts the overall BOLD activity during sleep-like states. The nucleus basalis located in the basal forebrain has widespread cholinergic projections to the neocortex and is an essential neuromodulator regulating arousal and attention (DiFrancesco MW *et al.* 2019). Hence, any withdrawal of excitatory cholinergic input from the basal forebrain can result in hyperpolarisation in the cortical pyramidal neurons leading to slow/large amplitude oscillations typical of deep sleep. The basal forebrain region may not only affect electrographic activity of the cortex via direct projections (Lin S-C et al.; Xu M *et al.* 2015) but also the cerebral blood flow in the basal forebrain-cingulate network resulting from the amount of intrinsic sleep-drive (Poudel GR *et al.* 2012). Our model therefore provides evidence for an important role of basal forebrain in seeding the spread of activation/deactivation during sleep-like intrusions in the awake brain.

## Notes

https://github.com/govin2000/inferringsleep

### Competing Interest Statement

The authors have declared no competing interest.

