## Supplementary Information for "Sleeping While Awake: The Intrusion of Neural Activity Associated with Sleep Onset in the Awake Human Brain"

**This PDF file includes:**

Supplementary text
Figures S1 to S5
Tables S1 to S2

**Other supplementary materials for this manuscript include the following:**

Data analysis scripts and datasets located at:
https://github.com/govin2000/inferringsleep

### Supplementary Text

#### Methods

*Experiment 1: Simultaneous fMRI and EEG study of light sleep*

Simultaneous EEG and fMRI data were acquired using a 64-channel MR compatible EEG system (BrainProducts, Germany) and a 3T scanner (Siemens, Skyra). The 12 participants spent approximately two hours in the lab. They were prepared with EEG head cap outside the MRI scanner room. Once each EEG electrodes were filled with conducting (Ag-Ag Cl) gels, the electrodes were tested for impedance via BraionVision Recorder software. Of the 64 channels available on the EEG cap (EasyCap, Brain Products, Germany), 62 channels on the head were positioned according to the International 10–20 system. An extra channel was used to record ECG. The electrode impedances were kept below 20 ohm for most electrodes. Once ready, the participants were taken to the MRI room for scanning. They laid supine on the MRI table with head coil foam-padded to restrict head motion and improve subject comfort. EEG data acquisition was synchronized with the MR scanner clock (Syncbox, Brain Products, Germany).

EEG data were processed to remove gradient and BCG artefacts using BrainVision Analyser and EEGLab software. The BrainVision Analyser software removed gradient and BCG artefacts from the data by using adaptive subtraction techniques implemented in the software. Any residual BCG artefacts were removed using independent component (ICA) analysis. To achieve this, the processed EEG data was imported into EEGLab software. ICA (using runica) was run on the EEG data within the EEGLab software. ICA separated

EEG into signal and noise components. Any components which were highly correlated (*r*>0.4) with ECG signal were removed from the data.

The fMRI data were processed using FSL (FMRIB's Software Library, www.fmrib.ox.ac.uk/fsl), ANTs (http://stnava.github.io/ANTs/), and custom Linux Shell and Matlab scripts (Matlab 7.6.0, R2008a, Mathworks, MA, USA). The fMRI data were preprocessed for motion correction (mcflirt), slice-time correction (FSL), and temporal filtering (256 s) using FSL FEAT. The pre-processed fMRI was normalised to a standard 2x2x2 mm MNI brain (MNI152) using linear and non-linear registration tools available in FSL and ANTS. The steps for normalisation included (1) Brain extraction of T1-weighted structural MRI using antsBrainExtraction.sh routine, (2) Segmentation of the brain into different tissues types using FAST routine available in FSL, (3) Linear registration of fMRI data to T1-weighted MRI using FLIRT routine (bbr registration using the tissues types obtained from FAST), (4) Non-linear registration of T1-weighted MRI to MNI152 2mm x 2mm x 2mm brain using antsRegistrationSyN.sh routine (5) Application of linear and non-linear transformations of the fMRI data into MNI space.

The preprocessed and normalised fMRI data were analysed using FSL FEAT to identify the fMRI activity associated with EEG theta and EEG alpha. The first-level model included (1) EEG Alpha, (2) EEG Theta, (3) 6 motion regressors from motion parameters (4) Large motion regressors from fsl_motion_outliers (with default dvars option), (5) the first eigenvector of fMRI activity within a CSF mask (to account for any physiological noise). The contrast parameter estimates from the first-level, measuring EEG-theta and EEG-alpha related fMRI activity, were analyzed using group-level non-parametric t-test (Randomise function in FSL).

**Experiment 2: fMRI study of logical decision making after partial sleep deprivation**

*Participants and protocol*

All 20 participants indicated a normal sleep schedule with a typical time to bed between 10:00pm–12:00pm and a usual time-in-bed of 7.0 to 8.5 h. Participants visited the laboratory on three occasions. During the first visit, they were informed about the full experimental procedure. Informed consent was obtained, and participants were given an Actiwatch (Respironics Inc., PA, USA) and detailed sleep diary to document their sleep habits. Participants were required to record this for 6 days and 5 nights prior to the two experimental sessions. They were requested to abstain from consumption of caffeine, nicotine, and alcohol for the duration of the experiment. The two subsequent visits followed a night of normal sleep (rested) or partial sleep deprivation. For the rested session, participants were directed to go to bed between 10:00pm–12:00pm and have a time-in-bed of 7.0 to 8.5 h. For sleep-deprived session they were to maintain normal sleep except for the night prior to the scanning day during which they were instructed to have only 4 hours time-in-bed.

*Game of set task*

The logical decision-making task used in the study is based on the rules for the game of 'Set'. In the game of Set, a Set consists of three cards in which each of four features (colour, symbol, number, and shading) is either the same on each card or is different on each card. Each card has a variation of the following four features:

(a) Colour: The symbols of each card are red, green, or purple.

(b) Symbol: Each card contains ovals, crosses, or diamonds.

(c) Number: Each card has one, two, or three symbols.

(d) Shading: Each card is solid, open, or striped.

For example, the following card is (a) green, (b) diamond, (c) three symbols, and (d) solid shading.

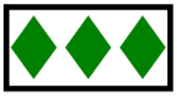

A 'Set' consists of three cards in which each feature is either the same on each card or is different on each card. For example, the following three cards are a Set because they *all* have the *same* colour, symbol, and number of symbols and *all* have *different* shading:

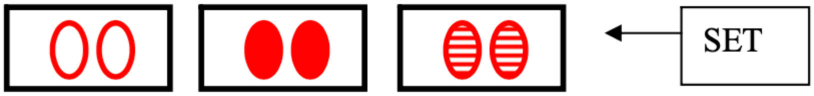

The following three cards are a Set because they *all* have the *same* number of symbols and shading and *all* have *different* symbols and colour.

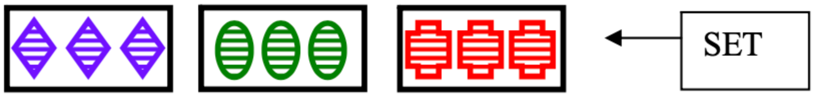

The following three cards are not a Set because – although they *all* have the *same* symbol and shading, and *all* have *different* number of symbols – two cards are purple and one is green.

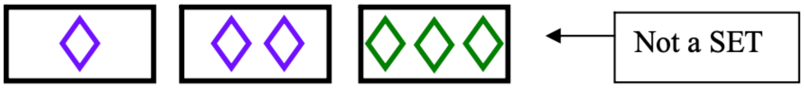

The *Golden Rule* is – if two cards are the same for any feature (colour, number, symbol, or shading) and one is not, then the three cards are not a Set.

In order to complete the task, they were presented with three cards on the computer screen. They had 5 s to decide whether the three cards are a Set or not and as quickly and accurately as you can press the left button 'Yes' or the right button for 'No' (Fig. S5).

**fMRI data collection and processing**

Participants in this study were scanned using a 3.0 T MRI Scanner (GE Medical Systems). The first five images from each session were disposed to permit T1 equilibration. fMRI data were acquired in all three runs of the decision-making task. The first four volumes were removed. Participants were provided with ear plugs to reduce the high-volume acoustic noise from the scanner. Additional pads were placed on both sides of the head to reduce head motion and further minimise acoustic noise.

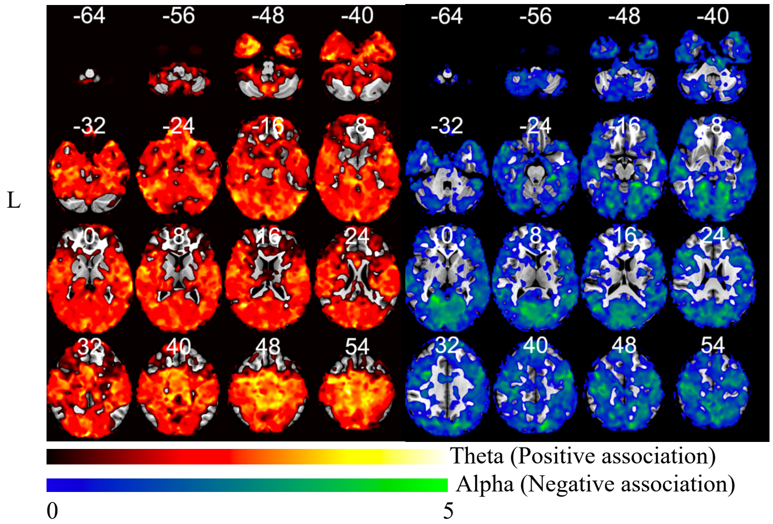

SI Fig. 1: Spatial distribution of t-statistics representing fMRI activity associated EEG theta (red-yellow) and EEG alpha (blue-green) power spectral density.

**Validation analysis of independent fMRI dataset from the Stockholm sleepy brains study**

The data from the sleepy brain project was downloaded from openfmri.org (accession number ds000201). Only fMRI scans with complete dataset (number of volumes = 200) and those that did not registration failured were used in the study (N=56). The fMRI data was preprocessed for slice-time correction (3dtshift), motion correction (mcflirt), co-registration with T1-weighted MRI scans (white matter segmentation), and normalization to standard space (using ANTS non-linear registration of T1 to MNI template). For each participant, first 5 eigenvectors of fMRI data within whitematter+CSF mask (compcorr), large motion outliers (identified using fsl_motion_outlier and default dvars option) were estimated. To ensure that our method is not dependent on a type of template choice, we used a different functional template with predefined set of 126 ROIs, which included 114 cortical regions derived from an independent analysis of whole-brain functional organization in a large sample of 1,000 subjects and 12 subcortical structures from the Automated Anatomical Labeling template[1].

The sleep-like events were inferred from the fMRI data using the approach described elsewhere. These events were used in the first-level analysis of fMRI data. The GLM used in the first-level analysis included inferred sleep-events and KSS sleepiness response as onsets. Confound regressors were 6 motion parameters and large motion outliers from fsl_motion_outlier (dvars with default option). The parameter estimates from the first-level analysis were used in a non-parametric group-level t-test (Randomise, 5000 permutations) to estimate overall transient activations across the individuals. Any

**Commented [NP1]:** Isn't it called open neuro now?

activations/deactivations significant at p<0.05 (family-wise corrected for voxels) were considered to be the co-activation of significance.

To estimate level of sleepiness during the scanning, we averaged Karolinska Sleepiness Scale (KSS) values during the 8 minutes. To determine whether sleepiness is associated with co-activation, we median split the average KSS scores into alert and sleepy individuals. An upaired-ttest was then performed on the first-level fMRI activity pattern associated with sleep-like events. Any brain regions showing significant difference (p<0.05, cluster corrected z>2.3) difference were visualised and reported.

(A) (B)

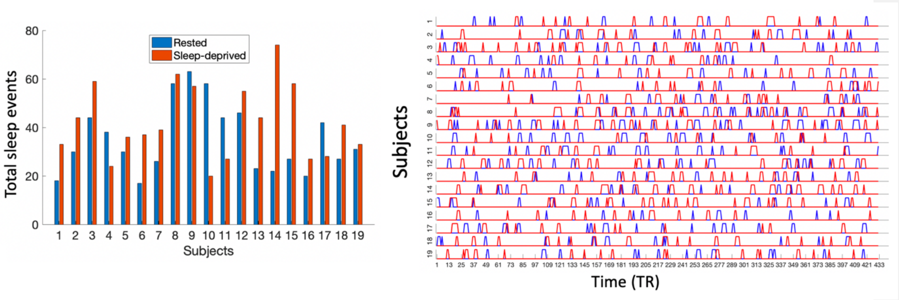

Fig S2: (A) Total number of inferred sleep events in the rested and sleep-deprived fMRI data. (B) Temporal location of sleep events across subjects in rested (blue) and sleep-deprived (red) sessions.

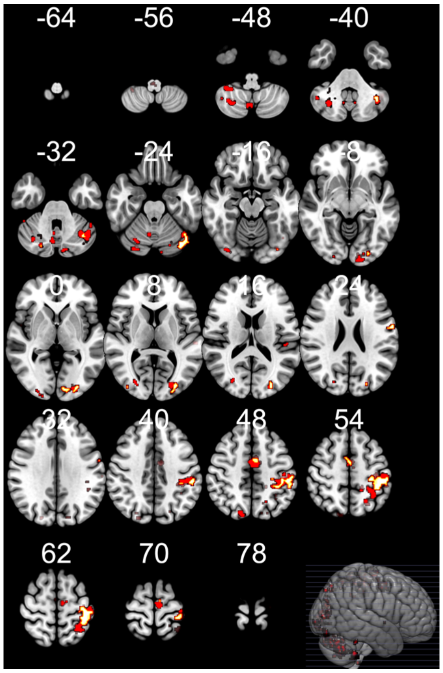

Fig S3: The brain regions showing overlap between task-related and sleep-related brain networks in rested (red) and sleep-deprived (hot) conditions.

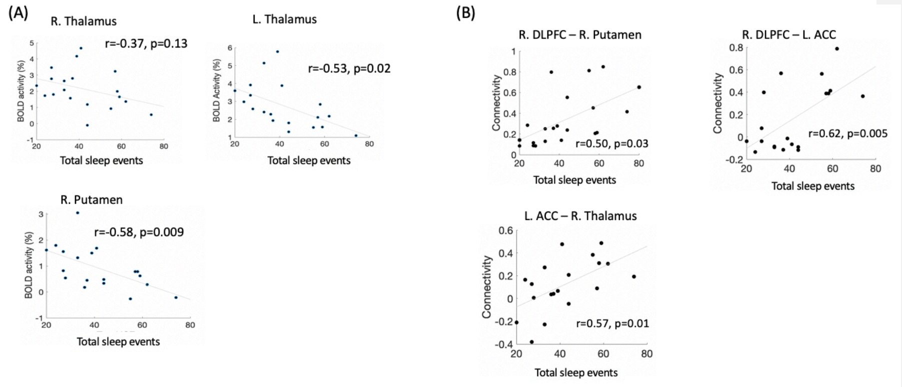

Fig. S4: Scatterplots of the correlation between task-related BOLD fMRI activity and total sleep events (A) and task-related fMRI connectivity and total sleep events (B).

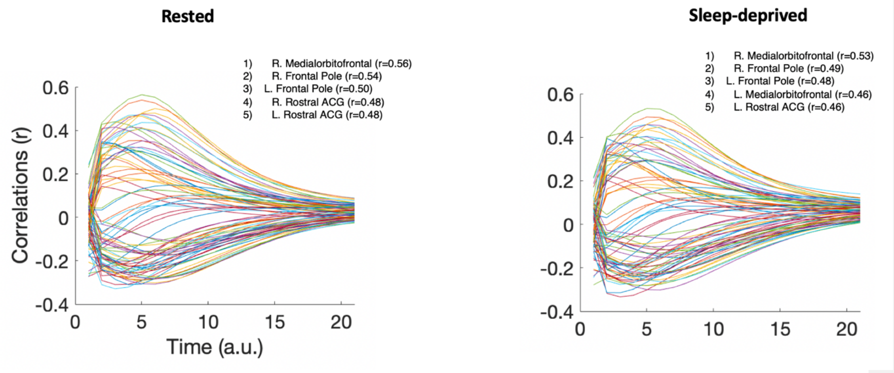

Fig S5: The correlate-time plots of the correlation between predicted and measured activity during onset of sleep. Each curves represent the similarity between predicted and measured activity over time for each of the 82 ROIs.

**Table S1:** The brain regions showing significant positive correlation with theta activity (3-7 Hz) and negative correlation with alpha (8-13 Hz) activity in EEG. The finding are significant after correcting for multiple comparison using cluster based thresholding (z threshold of 2.3, p<0.05). Separate clusters given below were identified by using 'cluster' command available in FSL toolbox. Only clusters with size greater than 5 voxels are reported.

| **Brain Region** | **Size (voxels)** | **T-stat (max)** | **MAX X (mm)** | **MAX Y (mm)** | **MAX Z (mm)** |
|---|---|---|---|---|---|
| ***Positive association with theta power*** | | | | | |
| L. Precuneus | 161 | 4.33 | -8 | -38 | 52 |
| R. Precuneus | 137 | 3.59 | 10 | -46 | 50 |
| L. Superior frontal gyrus | 100 | 3.65 | -14 | -10 | 76 |
| R. Postcentral gyrus | 52 | 4.78 | 2 | -40 | 72 |
| R. Precentral gyrus | 45 | 4.13 | 14 | -28 | 78 |
| L. Postcentral gyrus | 31 | 3.43 | -8 | -50 | 76 |
| R. Lateral occipital cortex | 20 | 3.72 | 16 | -60 | 70 |
| L. Lateral occipital cortex | 16 | 3.8 | -18 | -58 | 72 |
| L. Precentral gyrus | 11 | 3.3 | -2 | -30 | 56 |
| L. Superior parietal lobule | 10 | 3.03 | -12 | -56 | 62 |
| L. Superior frontal gyrus | 8 | 3.25 | -26 | -6 | 68 |
| R. Superior frontal gyrus | 8 | 3.71 | 24 | -4 | 72 |
| ***Negative association with alpha power*** | | | | | |
| R. Lingual gyrus | 5477 | 5.82 | 16 | -48 | -6 |
| L. Inferior temporal gyrus | 593 | 5.14 | -52 | -62 | -18 |
| R. Postcentral gyrus | 537 | 4.55 | 56 | -14 | 52 |
| R. Lateral occipital cortex | 302 | 5.67 | 52 | -70 | 18 |
| L. Middle temporal gyrus | 245 | 4.18 | -68 | -54 | 12 |
| R. Inferior frontal gyrus | 190 | 5.41 | 48 | 28 | 20 |
| R. Precuneus | 170 | 6.52 | 6 | -48 | 58 |
| R. Precentral gyrus | 163 | 4.52 | 32 | 8 | 32 |

| | | | | | |
|---|---|---|---|---|---|
| L. Precuneus | 126 | 4.93 | -12 | -52 | 54 |
| R. Middle frontal gyrus | 124 | 5.42 | 46 | 4 | 56 |
| L. Superior temporal gyrus | 70 | 4.64 | -66 | -14 | 4 |
| R. Postcentral gyrus | 68 | 4.12 | 6 | -36 | 58 |
| L. Parietal operculum cortex | 65 | 4.19 | -40 | -36 | 16 |
| L. Insular cortex | 54 | 5.13 | -40 | -12 | 6 |
| L. Occipital fusiform gyrus | 35 | 3.6 | -26 | -84 | -8 |
| L. Inferior temporal gyrus | 33 | 3.5 | -50 | -18 | -22 |
| R. Supramarginal gyrus | 26 | 4.32 | 44 | -36 | 36 |
| L. Superior parietal lobule | 23 | 3.7 | -24 | -46 | 50 |
| L. Lateral occipital cortex | 19 | 4.07 | -38 | -70 | 2 |
| L. Lateral occipital cortex | 16 | 3.57 | -32 | -68 | 28 |
| R. Parahippocampal cortex | 15 | 5.06 | 28 | -30 | -26 |
| L. Angular gyrus | 8 | 3.39 | -50 | -54 | 34 |
| R. Supracalcarine cortex | 8 | 3.13 | 6 | -86 | -12 |
| R. Temporal occipital fusiform gyrus | 7 | 3.7 | 34 | -44 | -16 |
| L. Superior parietal lobule | 5 | 2.98 | -22 | -54 | 68 |
| L. Middle temporal gyrus | 5 | 3.45 | -48 | -40 | -8 |

**Table S2:** Significant task-related ($p<0.05$, voxel wise FEW corrected) fMRI activity in the brain during well-rested sessions. The coordinates represent local maxima of large clusters encompassing multiple brain regions.

| Brain Region | Z-value | X | Y | Z |
|---|---|---|---|---|
| L. Superior Lateral Occipital Cortex | 13.8 | -26 | -78 | 32 |
| L. Pallidum | 12.7 | -14 | -16 | -8 |
| R. Insula | 9.63 | 34 | 20 | -10 |
| R. Precentral Gyrus | 9.3 | 30 | -8 | 60 |
| R. Precentral Gyrus | 8.33 | 50 | 2 | 30 |
| L. Precentral Gyrus | 7.84 | -40 | -4 | 32 |
| R. Middle Frontal Gyrus | 7.64 | 42 | 26 | 24 |
| Brainstem | 7.77 | -4 | -44 | -68 |
| L. Parietal Operculum Cortex | 7.62 | -46 | -26 | 14 |
| L. Posterior Cingulate Gyrus | 7.14 | -6 | -38 | 24 |
| L. Middle Frontal Gyrus | 7.47 | -36 | 18 | 30 |
| L.. Postcentral Gyrus | 8.1 | -16 | -42 | 54 |
| R. Caudate | 6.55 | 14 | 12 | -2 |
| L. Inferior Frontal Gyrus | 7.74 | -50 | 32 | 22 |
| L. Frontal Pole | 6.91 | -34 | 48 | 22 |
| L. Anterior Cingulate Gyrus | 6.68 | -6 | 2 | 28 |
| R. Juxtapositional Lobule Cortex | 7.21 | 2 | 0 | 70 |
| L. Middle Frontal Gyrus | 6.65 | -42 | 8 | 30 |

1 Yeo, B. T. *et al.* The organization of the human cerebral cortex estimated by intrinsic functional connectivity. *J Neurophysiol* **106**, 1125-1165, doi:10.1152/jn.00338.2011 (2011).
